# Early Postnatal Tobacco Smoke Exposure Aggravates Experimental Autoimmune Encephalomyelitis in Adult Rats

**DOI:** 10.1101/2020.07.28.224709

**Authors:** Zhaowei Wang, Liping Wang, Fangfang Zhong, Chenglong Wu, Sheng-Tao Hou

## Abstract

Although substantial evidence supports smoking as a risk factor for the development of multiple sclerosis (MS) in adulthood, it remains controversial as to whether early-life exposure to environmental tobacco smoke (ETS) increases the risk of MS later in life. Here, using experimental autoimmune encephalomyelitis (EAE) as an animal model for MS, we show that exposing neonatal rats during the 1^st^ week (ETS1-EAE), but not the 2^nd^ week (ETS2-EAE) and the 3^rd^ week (ETS3-EAE) after birth, increased the severity of EAE in adulthood in comparison to pups exposed to filtered compressed air (AIR-EAE). The EST1-EAE rats showed a worse neurological deficit score and a significant increase in CD4^+^ cell infiltration, demyelination, and axonal injury in the spinal cord compared to AIR-EAE, ETS2-EAE, and ETS3-EAE groups. Flow cytometry analysis showed that the ETS1 group had decreased numbers of regulatory T (Treg) cells and increased effector T (Teff) cells in the brain and spinal cord. The expressions of Treg upstream regulator Foxp3 and downstream cytokines such as IL-10 were also altered accordingly. Together, these findings demonstrate that neonatal ETS exposure suppresses Treg functions and aggravates the severity of EAE, confirming early-life exposure to EST as a potential risk factor for multiple sclerosis in adulthood.

## INTRODUCTION

Multiple sclerosis (MS) is an autoimmune disease of the central nervous system, and its etiology is related to genetic and environmental factors (Thompson et al., 2018). Tobacco smoke has received increasing attention as a potential environmental risk factor for MS (Öberg et al., 2011; Carreras et al., 2019). Adult population case-control studies have shown that smoking or exposure to environmental tobacco smoke (ETS) promotes the disease progression and increases the recurrence of MS (Alphonsus and D’Arcy, 2019; Rosso and Chitnis, 2020). However, evidence for the impact of early-life exposure to ETS on adulthood MS development is not certain and even conflicting in the literature (Montgomery et al., 2008; Pantazou et al., 2015; Pronovost and Hsiao, 2019).

For example, a Swedish population-based register that prospectively recorded smoking during pregnancy assessed the impact of maternal smoking during pregnancy on the risk of developing MS later in life. The study found that prenatal smoking was not associated with the onset of MS amongst the offspring (Montgomery et al., 2008). Another case-control study involving 238,381 female nurses who self-reported their prenatal and perinatal factors also found that there was no correlation between prenatal and childhood exposure to ETS and MS risk (Gardener et al., 2009).

However, strong evidence also exists, supporting a positive correlation between early-life ETS exposure and the development of MS. It appears that early exposure to ETS within the first year of birth represents a critical time window. Case-control studies on the risk factors for the onset of MS in children revealed that the incidence of MS in patients exposed to tobacco smoke in the first year after birth had tripled (Graves et al., 2017). Moreover, maternal prenatal smoking was also positively associated with MS hospitalization later in life (Mueller et al., 2013). Nevertheless, one caveat of these studies is that a causal relationship can’t be drawn solely based on human population studies since the risk factors of developing MS later in life is multifaceted and may not be simply due to early-life ETS exposure. New findings in animal studies are emerging strengthening the idea that exposure to tobacco smoke in a critical period interferes with brain development of mice from late infancy to early adulthood (Wu et al., 2009, 2012; Torres et al., 2019, 2020).

CNS inflammation is one of the most common features of MS. The regulatory T (Treg) cells play a crucial role in maintaining immune homeostasis, and their limited differentiation or dysfunction is closely related to the development of MS (Josefowicz et al., 2012; Ohkura et al., 2013). Foxp3 plays a significant role in the development and function of Treg cells (Sakaguchi et al., 2010). The genetic defects of Foxp3 will cause the dysfunction of Treg cells and autoimmune diseases (Nie et al., 2015).

Based on these understandings, induction of experimental autoimmune encephalomyelitis (EAE) in rats was used as an animal model for MS to determine the impact of early-life ETS exposure as a risk factor for developing EAE in adulthood. Three specific time windows in early-life ETS exposure were also designed to test their relationship with EAE development in adulthood. We hypothesized that early-life ETS exposure has a long-term impact on the immune system and may aggravate the severity of MS. These animal model studies will provide evidence to overcome the limitations of the retrospective clinical research and ethical restrictions of testing on human subjects.

## MATERIALS AND METHODS

### Animals and ETS treatment

The animal experiments involved in this study were carried out in strict accordance with the Zhejiang Province Laboratory Animal Management Measures, and the local animal care committee approved the protocol. Sprague–Dawley pregnant rats were obtained from Shanghai Laboratory Animals Center (Shanghai SLAC Laboratory Animal Co., Ltd, Shanghai, China) and housed under controlled conditions (14 h light/10 h dark cycle with lights on at 7 AM, room temperature at 22 ± 2 °C).

Littermates at postpartum day 1 (P1) were separated into groups to receive ETS exposure as described previously (Teague et al., 1994; Slotkin et al., 2001). Briefly, to simulate the ETS exposure, sidestream smoke was generated using smokes from commercial cigarettes (Zhonghua™, 11 mg of tar, and 1 mg of nicotine per cigarette, Shanghai Cigarette Factory, Shanghai, China) and sent into a conditioning chamber for 2 min before dilution with filtered air. Whole dams were exposed to the side stream smokes in the exposure chamber for 1 h each time, twice per day. Three groups of pups were used for ETS treatment: ETS1 from P1 to P7, ETS2 from P8 to P14, and ETS3 from P15 to P21. During the period of passive smoke exposure, the nicotine concentrations, suspended particulate and carbon monoxide were measured at 108 ± 31 µg/m^3^, 1.21 ± 0.12 mg/m^3^, and 6.2 ± 0.32 ppm, respectively. The smoke total particulate matter concentration inside the exposure chamber was 220 ± 28 mg/m^3^. Control pups were exposed to filtered compressed air for 1 h each time, twice per day, from P1 to P21 (AIR group). All rat pups remained with their dams until they were weaned at 21 days of age.

### EAE Induction

EAE was induced as previously described at 10 weeks of age (Wang et al., 2011). Briefly, fresh guinea pig spinal cord homogenate was mixed with complete Freund’s adjuvant (containing BCG 10 mg/mL) in equal volumes, and repeatedly whipped in an ice-water mixture with a syringe until a water-in-oil emulsion was formed. The emulsion mentioned above was injected subcutaneously on the limb pads of rats at a total volume of 400 µl. The control group was injected with an emulsion of 0.9% sodium chloride solution mixed with complete Freund’s adjuvant serving as a control (CTL). The rats were then injected i.p. with 0.1 ml pertussis vaccine suspension (Shanghai Institute of Biological Products, China) at 0 and 48 h postimmunization.

### Evaluation of neurobehavioral deficits

The neurobehavioral deficits score was assessed from the first day after EAE induction until all the animals were sacrificed. The scoring method used for neurobehavioral defects was as previously described (Wang et al., 2009, 2011): grade 0, no obvious movement difficulties; grade 1, tail paralysis; grade 2, paresis of hind legs; grade 3, complete paralysis of hind legs; grade 4, tetraplegia; and grade 5, moribund state or death.

### Histology and immunohistochemistry

After 22 d of EAE induction, rats were anesthetized with 10% chloral hydrate and then sacrificed by infusing the left ventricle with 0.9% sodium chloride solution. The spinal cord tissue was isolated and fixed in 40 g/L paraformaldehyde for 24 h, and then dehydrated, cleared, and embedded in wax to make wax blocks. Paraffin sections of 5 µm thickness were cut and stained with hematoxylin-eosin (H&E), Luxol fast blue staining (LFB), and silver glycine staining according to standard methods to determine the histopathology EAE damage to the spinal cord. The procedure of silver glycine staining was performed using a commercial kit according to the manufacturer’s instructions (Servicebio, Wuhan, China).

Rabbit anti-CD4, anti-GFAP polyclonal antibodies (Abcam, USA) were used to detect CD4^+^ T cells and astrocytes at a dilution of 1:150 and 1:200, respectively. The method of immunostaining is the same as previously described (Hou et al., 2006; Yu et al., 2019; Zheng et al., 2020). Briefly, after treatment with 0.03% H2O2 to inactivate endogenous peroxidase, the concealment of antigen was achieved using microwaves. Non-specific binding was blocked with 10% fetal bovine serum (GIBCO, Australia), and the sections were incubated with the primary antibody overnight at 4°C. After washing with PBS, sections were incubated with biotinylated goat anti-rabbit antibody (1:100). An avidin-biotin complex kit was used to visualize staining (VECTASTAIN ABC Kits, Vector Laboratories, USA).

### Tissue section quantifications

Scoring of H&E staining was according to the method described previously (Okuda et al., 1999; Wang et al., 2009, 2011) with a total score of 4 points; 0 points: no inflammatory changes; 1 point: inflammatory cell infiltration is limited to the perivascular and meningeal; 2 points: mild inflammatory cell infiltration in the spinal cord 1 - 10 cells/piece); 3 points: moderate infiltration of inflammatory cells in the spinal cord (11 - 100 cells/piece); 4 points: severe infiltration of inflammatory cells in the spinal cord (> 100 cells/piece). Axonal damage was semi-quantitated through randomly selected fields from the white matter in each spinal cord area to provide digital images. The number of axons per square millimeter counted and plotted as previously described (Ellestad et al., 2009). Quantitative analysis of CD4 cell counts per square millimeter was performed as previously described (Wang et al., 2011). From three randomly selected high-power fields of view at ×400 times of magnifications under a microscope, a scanner was used to count the number of infiltrating CD4^+^ cells. Image J (National Institutes of Health, Bethesda, MD) was used to quantify the level of LFB intensity. A double-blind approach was used in all quantitative studies.

### Flow cytometry

Cells were surface stained with fluorescent dye-conjugated antibodies that recognize CD25 and CD4. CD4 and CD25 antibodies were obtained from Multisciences (LIANKE) Biotech Co. (Hangzhou, China). Spleen cells were separated, homogenized and harvested after washing with PBS and staining with surface markers. BD FACSDiva™ software was used to analyze the stained cells on BD Fanscan to flow cytometry for collection. Flow cytometry data were analyzed by using FlowJo V10.6.2 according to previously published methods (Wang et al., 2011)

### Real-time RT-PCR

The monocytes obtained the brain and spinal cord homogenate were isolated and purified by Percoll density gradient centrifugation. The CD4^+^ cells from the cerebrospinal cord were obtained using flow cytometry, as previously described (Du et al., 2009). CD4^+^cells were homogenized in TRIzol (Invitrogen, USA), and the total RNA was purified from the aqueous phase. After RNA was treated with DNase I (Promega, USA) according to the scheme recommended by the manufacturer, cDNA was prepared by using MMLV reverse transcriptase (epicenter) to start reverse transcription of RNA by Oligo (dT) 18 (Shanghai Biotechnology Co, Shanghai, China) Real-time PCR was carried out on the Rotor gene 3000 real-time PCR (Corbett Research, USA). All PCR primers used in this article are provided in Table 1. All PCR data were normalized to GAPDH mRNA levels.

**Table 1.**
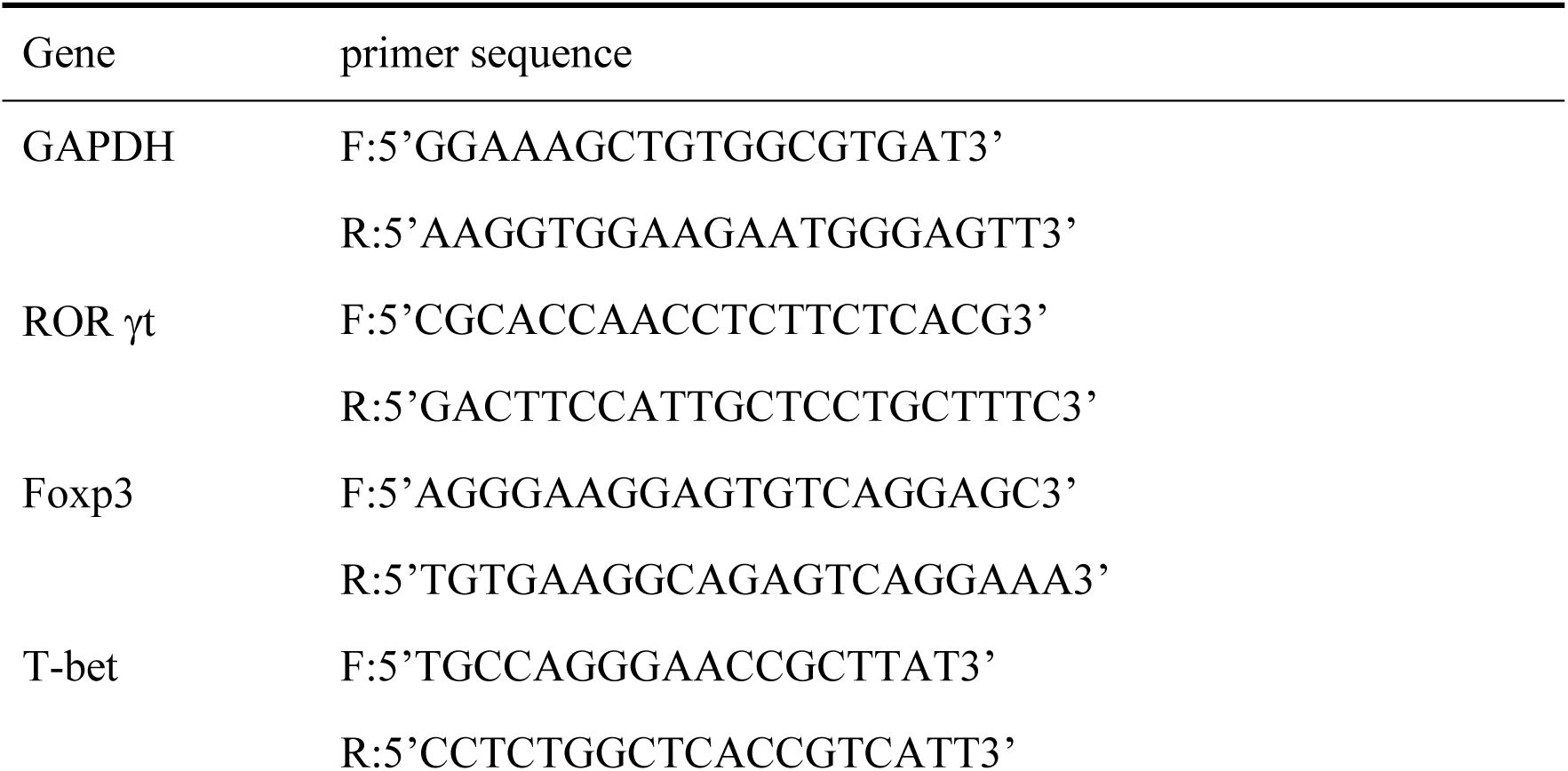
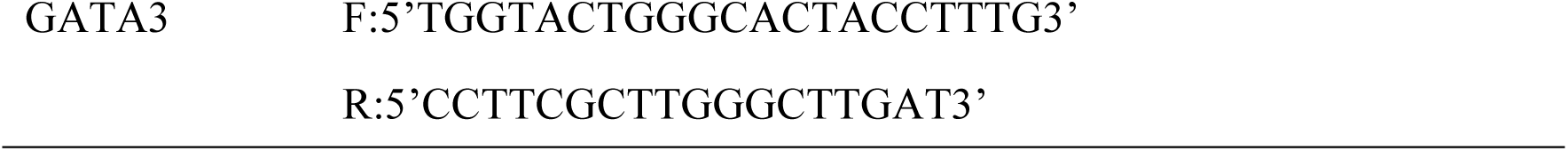
PCR primers.

### ELISA

The rats were anesthetized and fixed on the stereotactic frame. Cerebrospinal fluid (CSF) was obtained from the cistern by butterfly needle and syringe. CSF samples were rapidly frozen in liquid nitrogen and stored at −70°C until treatment. CSF samples were thawed, and the concentrations of IL-17F, IFN-γ, IL-4, and IL-10 were detected using the method provided by the kit manufacturer (R&D Systems, USA). Optical densities were measured using a Model 680 microplate reader (Bio-RAD, Hercules, CA) at 450 nm.

### Statistical analysis

All analyses were performed using Prism (Version 7) software. Unpaired t-test, one-way ANOVA or repeated two-way ANOVA were used with appropriate post hoc analysis as indicated in the text to identify significant groups. Spearman rank correlation coefficient-test was used to determine the correlation of nonparametric distribution data, and the linear regression analysis was performed using GraphPad Prism. Data are shown as mean ± SEM, with *P* > 0.05 considered not significant (n.s.). A *P* value < 0.05 was taken to indicate statistical significance. The symbol *indicates *P* < 0.05; ** indicates *P* < 0.01; and *** indicates *P* < 0.001.

## RESULTS

### Early postnatal ETS exposure aggravates the severity of EAE in adulthood

Rat littermates were randomly separated into three groups to receive exposure to tobacco smoke from P1 to P7 (EST1-EAE group, n = 18), P8 to P14 (EST2-EAE group, n = 18), and P15 to P21 (EST-EAE3 group, n = 18). Their mothers were also exposed to the same tobacco smoke treatment at the same time. Rat littermates exposing to filtered air were used as controls (AIR-CTL, AIR-EAE groups) (Fig. 1a). At the age of 10 weeks, EAE was induced in rats, and the AIR-CTL and EST-CTL groups received CFA serving as a control for EAE. The neurobehavioral deficits were measured at the beginning of the induction of EAE every day for 22 days, as shown in Fig. 1a.

**Fig. 1.**
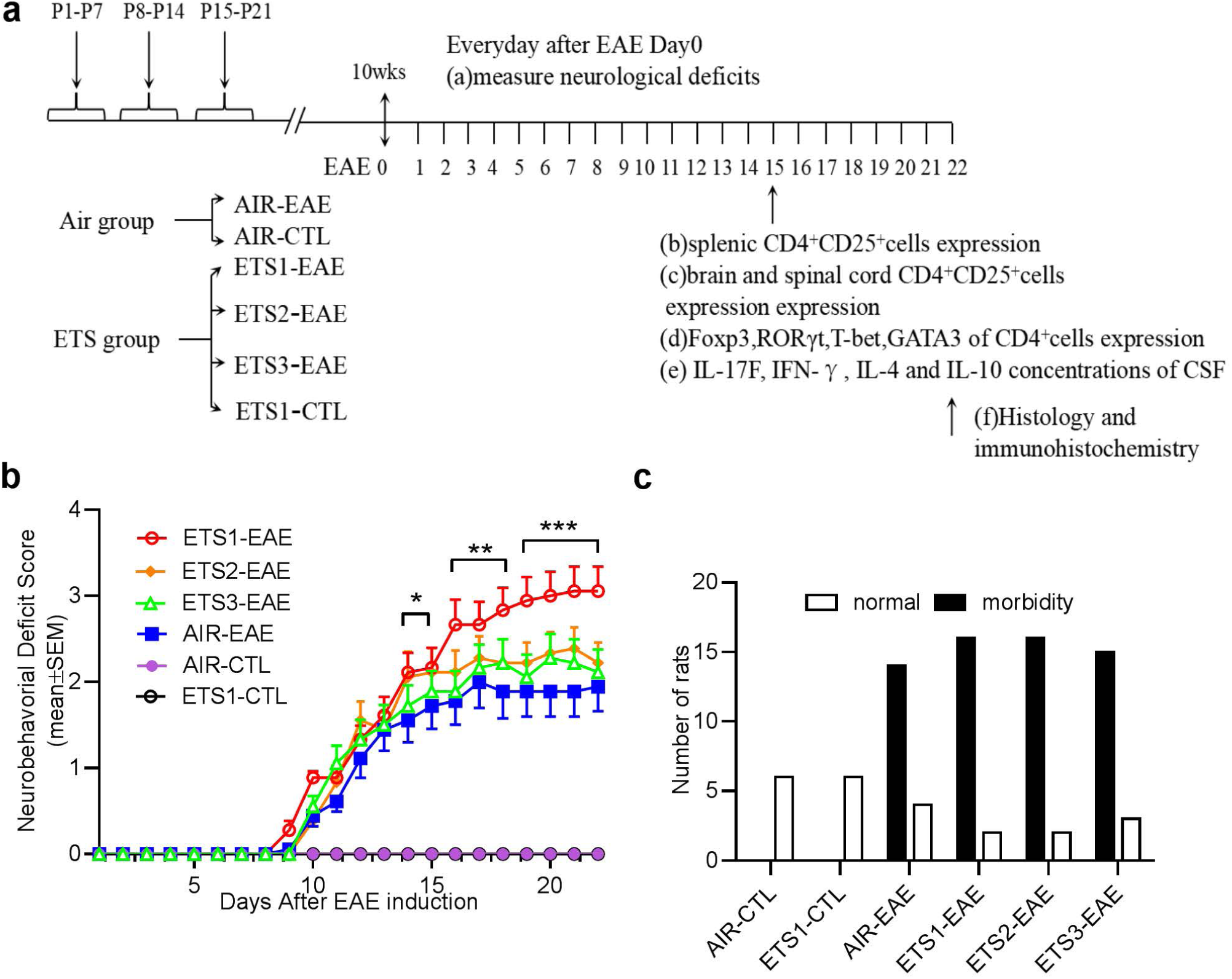
Early postnatal exposure to ETS aggravates EAE severity in adulthood. (a), A schematic diagram showing the time when ETS was given to the pups. EAE was induced in 10 wks old rat littermates. The EAE induction day was considered as day 0 for EAE development. Vertical arrows indicate the time when biological sampling occurred. (b), Neurobehavioral deficits were measured every day since EAE induction at day 0 (n = 18 in each group), and plotted in panel b. (c), The occurrence rate of EAE (Morbidity) was at 0 in the two CTL groups. No statistical significance occurred amongst all EAE induction groups, as determined by the chi-square test. Data represent the mean ± SEM. Error bars indicate SEM. n = 18 mice in EST groups and n = 6 in the AIR groups. * *P* < 0.05; Test used in panel b: repeated tow-way ANOVA with Sidak’s post hoc analysis;

Early postnatal exposure to tobacco smoke significantly aggravates the severity of EAE in adult littermates. As shown in Fig. 1b, the ETS1-EAE group had a significantly higher neurobehavioral deficit score compared with the AIR-EAE group at day 9 - day 22 (Fig. 1b, repeated two-way ANOVA with Sidak’s post hoc test at \**P <* 0.05, \*\**P* < 0.01, \*\*\**P* < 0.001; n = 18), and in contrast, there were no significant differences compared with the ETS2-EAE group (*P* = 0.6962, n = 18), and ETS3-EAE group (*P* = 0.8971, n = 18), respectively (Fig. 1b). There were no neurobehavioral deficits in the AIR-CTL group and the ETS1-CTL group. Furthermore, as shown in Fig. 1c, the occurrence rate of EAE (morbidity) was at 0 in the two CTL groups, while there was no statistical significance amongst all EAE induction groups as determined by the chi-square test (χ^2^ = 1.180, DF = 5, *P* = 0.9468). Together, these data demonstrated that early postnatal exposure to tobacco smoke significantly increased the severity of neurological deficits of EAE in adult rats.

### Early postanal ETS exposure aggravates inflammation of the spinal cord in EAE rats

To demonstrate the histopathological effect of ETS exposure on EAE severity, rat spinal cord tissue was extracted 22 days after EAE and sectioned for staining with H&E, LFB, CD4, and GFAP (Fig. 2a). EAE is an autoimmune disease dominated by cellular immunity. The most characteristic feature of EAE is the infiltration of inflammatory cells along the spinal cord membrane, and the severity can be manifested as the patchy aggregation of lymphocytes distributed along the small blood vessels (Fig. 2a). H&E staining of the cervical spinal cord cross-section revealed no infiltration of inflammatory cells in all CFA-treated control rats. Semi-quantitative analysis of H&E staining showed that the ETS1-EAE group had a higher lymphocyte infiltration score than that of the AIR-EAE group (Fig. 2b, *P* = 0.0238 by one-way ANOVA with Dunnett’s post hoc test, n = 18). Comparisons between ETS2-EAE and ETS3-EAE groups with AIR-EAE group, respectively, in H&E lymphocyte infiltration scores showed no statistical significance (Fig. 2b, one-way ANOVA with Dunnett’s post hoc test, AIR-EAE vs. ETS1-EAE, *P* = 0.0238; AIR-EAE vs. ETS2-EAE, *P* = 0.1392; AIR-EAE vs. ETS3-EAE, *P* = 0.6316; n = 18).

**Fig. 2.**
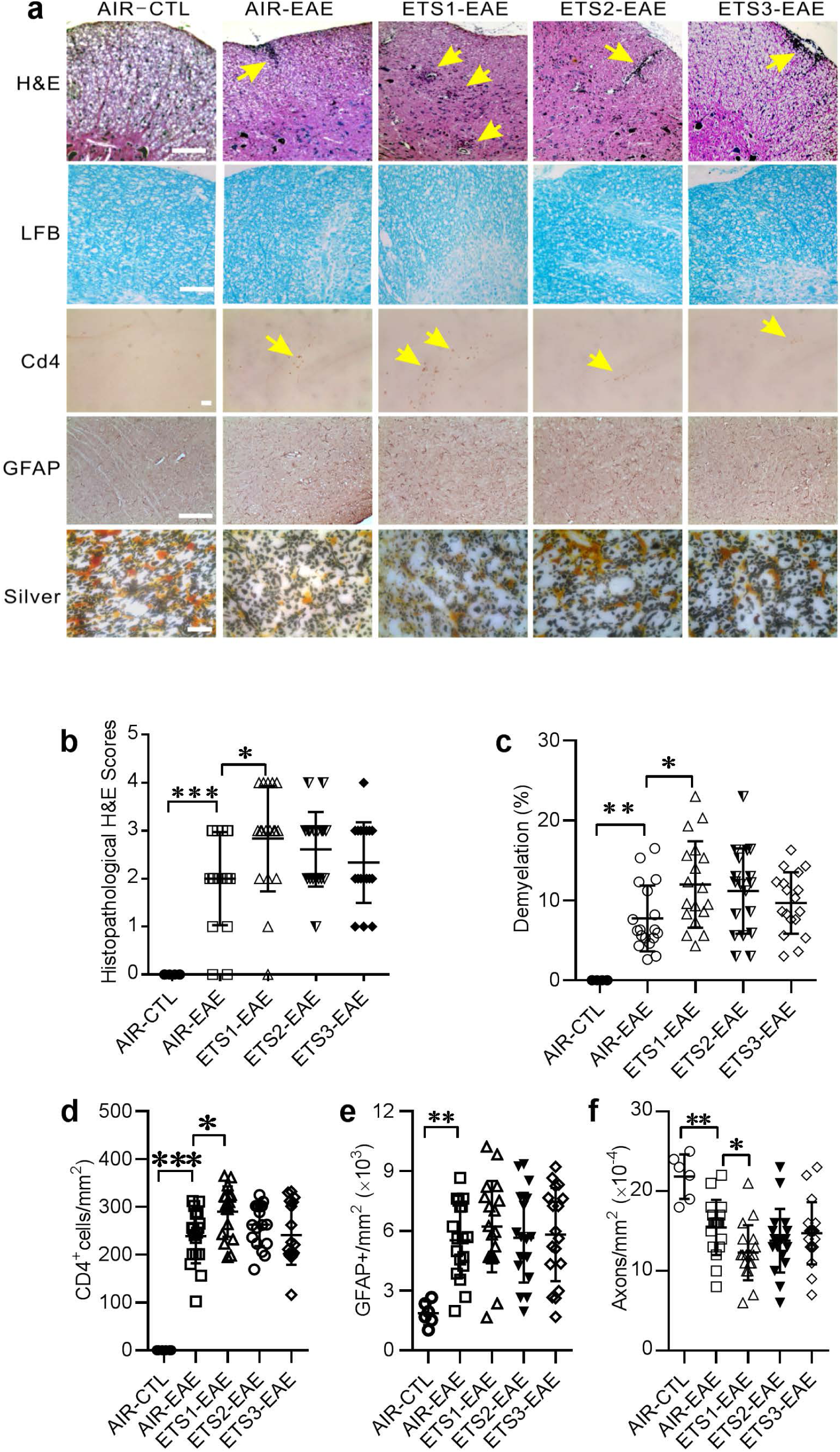
Histochemical evidence of spinal cord inflammation in the EAE rats. The rat cervical spinal cord was isolated 22 days after EAE induction. Spinal cord tissue was sectioned for staining of H&E, LFB, and silver. Immunohistochemical staining of CD4 and GFAP were also performed, as shown in panel (a). Arrows indicate perivascular inflammatory cell infiltration in the spinal cord under H&E staining. Sever demyelination (white color) was shown in the second row with LFB staining. In the third row, CD4^+^cells were stained brown using immunohistochemistry (arrows). Because of the scratches on the microscope lens, a similar pattern of background is visible in every micrograph in the third row of CD^4+^ staining. In the fourth row, GFAP immunohistochemical staining showed that astrocytes were stained brown. The fifth row was silver-stained axons shown in black. (b), Semi-quantitative analysis of H&E staining scoring. (c), Semi-quantitative analysis of the percentage of the demyelinating area of LFB staining. (d), The infiltration CD4^+^ cells counted in spinal cord tissue per unit area under the light microscope. (e), Comparison of axon density in the white matter area of the cervical spinal cord in each group. Scale bars = 50 µm. Data represent the mean ± SEM. Error bars indicate SEM. n = 18 in each EAE group and n = 6 in CTL group. * *P* < 0.05, ** *P* < 0.01; Test used: one-way ANOVA with Dunnett’s post hoc analysis.

Loss of myelination in the spinal cord was determined using LFB staining. CFA-treated rats showed no significant loss of myelin in the white matter area of the spinal cord (Fig 2a). The demyelination of the ETS1-EAE group was significantly severer than that of the AIR-EAE group by a semi-quantitative analysis (Fig. 2c, *P* < 0.05, n = 18). Compared with the AIR-EAE group, the demyelination of ETS2-EAE and ETS3-EAE groups were more serious, however, not statistically significant by the semi-quantitative analysis (Fig. 2c, n.s. *P* > 0.05, n = 18).

Central infiltration of CD4^+^ cells also reflects the severity of the inflammatory response. Using CD4 antibody immunohistochemical staining revealed that the ETS1-EAE group, compared with the AIR-EAE group, had a significantly increased number of CD4^+^ cells infiltrated in the cervical white matter area (Fig. 2a and d, *P* < 0.05, n = 18). In contrast, compared with the AIR-EAE group, infiltration of CD4^+^ cells in the ETS2-EAE and ETS3-EAE groups was not significantly different, albeit at a higher level (Fig. 2a and d, *P* < 0.05, n = 18).

Severe demyelination during the pathogenesis of EAE causes axon damage leading to the deterioration of nerve functions. Techniques of silver staining on paraffin sections were used to show the loss of axons in EAE spinal cords. CFA treatment did not cause damage to axons, while EAE rats had significantly lower axonal density than that of the CFA-treated rats (Fig. 2a and e, *P* < 0.001). Specifically, compared with AIR-CTL, the axon density in AIR-EAE groups decreased significantly (Fig. 2a and e, *P* = 0.0015, n = 6 in AIR-CTL, and n = 18 in AIR-EAE). Interestingly, the EST1-EAE has a significantly lower axonal density compared with the AIR-EAE group, demonstrating a worse outcome in rats exposed to tobacco smoke early in life (Fig. 2a and e, *P* = 0.0388, n = 18).

Increased GFAP positive astrocytosis is an indicator of a robust inflammatory response. There was no apparent glial proliferation in CFA-treated rats. Elevated GFAP cells occurred in all EAE rats (Fig 2a; one-way ANOVA with Dunnett’s post hoc test, *P* = 0.0018). However, there was no significant difference in the degree of astrocytosis between the AIR-EAE group and the all three ETS-EAE groups (Fig. 2a; one-way ANOVA with Dunnett’s post hoc test: AIR-EAE vs. ETS1-EAE, *P* = 0.7335; AIR-EAE vs. ETS2-EAE, *P* = 0.9991; AIR-EAE vs. ETS3-EAE, *P* = 0.9822).

Collectively, these results demonstrated that postnatal days 1-7 represent a critical time window for the development of sensitivity to EAE later in life in response to environmental tobacco smokes.

### Early-life ETS exposure affects the differentiation of central infiltrating T cells

Treg cells (CD4^+^CD25^+^) play an important role in inhibiting the hyperactivity of the immune system and promoting the recovery of autoimmune disease, while the effector T cells (Teff, CD4^+^CD25^-^ cells) participate mainly in the process of disease development. Results of quantitative analysis of the numbers of CD4^+^ cells infiltrating the spinal cord tissues using flow cytometry is consistent with the result obtained using CD4 immunohistochemical staining shown in Fig. 2a and d (Fig. 3a and d, *P* = 0.0070, n = 8).

**Fig. 3.**
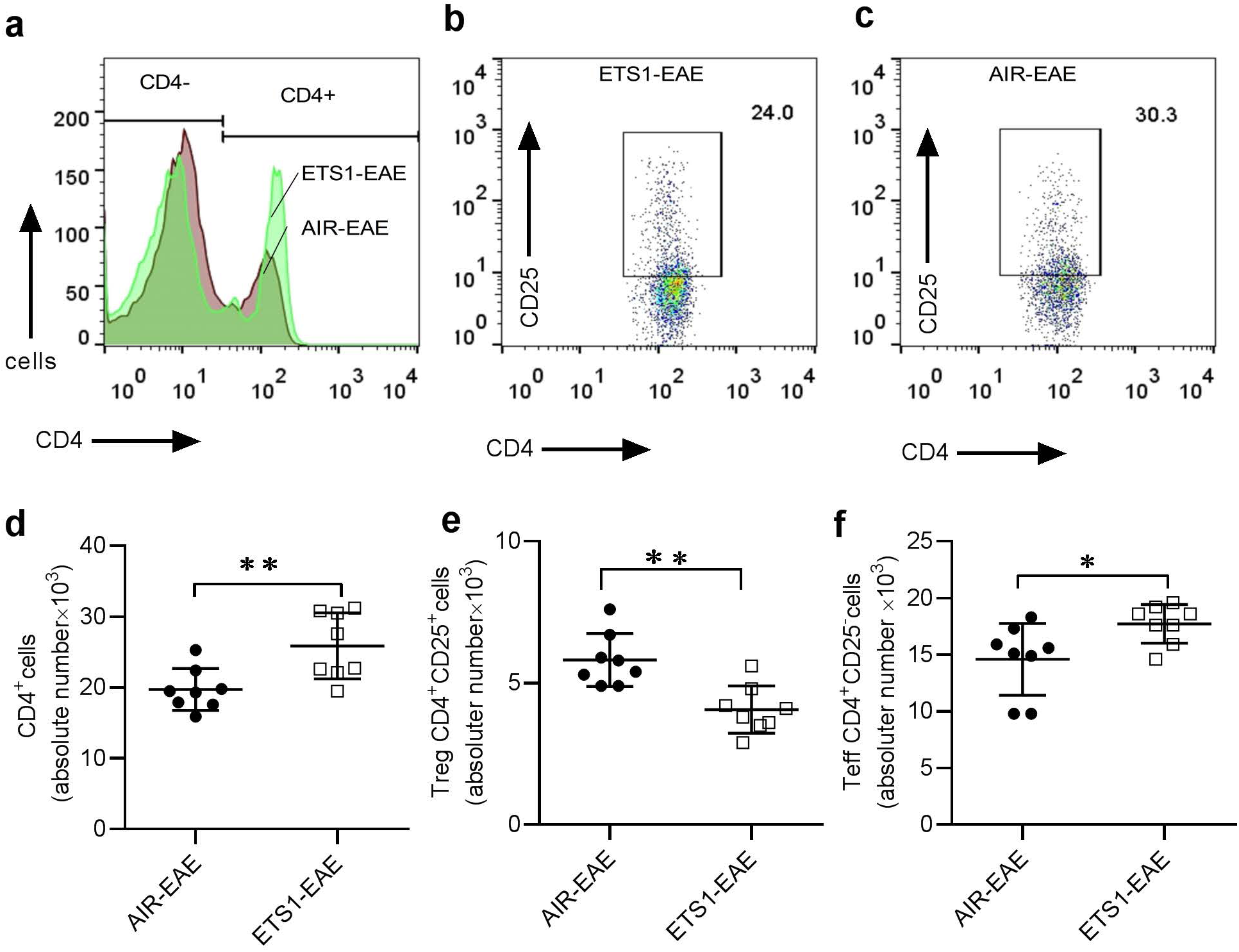
Effects of ETS exposure on the differentiation of CD4^+^cell subtypes. The monocytes from the brain and spinal cord were isolated and purified after the rats were killed on the 15^th^ day after EAE induction. CD4 and CD25 fluorescent-labeled antibodies were used to detect peripheral and central lymphocyte subtypes by flow cytometry. (a) Quantitative analysis of the numbers of CD4^+^ T cells derived from the monocytes infiltrating the brain and spinal cord tissues. (b,c), Quantitative analysis of infiltrating monocyte subtypes by CD4 and CD25 cell surface staining. The data displayed in the square gate indicates the percentage of cells in the CD4^+^CD25^+^ gate. (d), The absolute numbers of CD4^+^CD25^+^ Treg cells in monocytes from the brain and spinal cord (\*\**P* < 0.01, n = 8). (e), The absolute numbers of CD4^+^CD25^-^ Teff cells in monocytes from the brain and spinal cord (\**P* < 0.05). Data represent the mean ± SEM. n = 8 rats in both EAE groups. * *P* < 0.05, ** *P* < 0.01; Test used: Unpaired t-test analysis.

To determine the effect of early-life ETS exposure on the differentiation of Treg and Teff, monocytes were isolated from the homogenates of the spleen, brain, and spinal cord of rats at the 15 d induction of EAE. ETS exposure did not change the proportion of Treg and Teff cells in the spleen (n = 8, *P* > 0.05). In contrast, further analysis of lymphocytes in the brain and spinal cord showed that ETS exposure reduced the numbers of Treg (Fig. 3b and e, *P* = 0.0014, n = 8) and increased Teff (Fig. 3b and f, *P* = 0.0276, n = 8). These data indicated a possible effect of early-life ETS might inhibit Treg recruitment to the CNS and promote Teff cell invasion to the CNS.

### Early-life ETS exposure down-regulates Foxp3 expression and is related to the deterioration of EAE

Given the present findings of the down-regulation of Treg in the brain and spinal cords of early-life ETS exposed animals, we examined the FoxP3 transcription level of CD4^+^cells in the brain and spinal cord. At the same time, we also detected the mRNA expression levels of the other three transcription factors, RORgt, T-bet, and GATA3, which represent the activity of Th17 cells, Th1 cells, and Th2 cells, respectively. Compared with the AIR-EAE group, the expression level of Foxp3 in CD4^+^ cells of the ETS1-EAE group was significantly lower (*P* = 0.0312, n = 8). However, no differences occurred in RORgt, GATA3, and T-bet (Fig. 4a). Using a correlation analysis (Fig. 4b), we found that the expression level of Foxp3 (R^2^ = 0.2747, r = −0.5241), but not RORγt (R^2^ = 0.0098), GATA3 (R^2^ = 0.0179), and T-bet (R^2^ = 0.1130), was negatively correlated with the neurobehavioral deficit scores (Fig. 4b, n = 16).

**Fig. 4.**
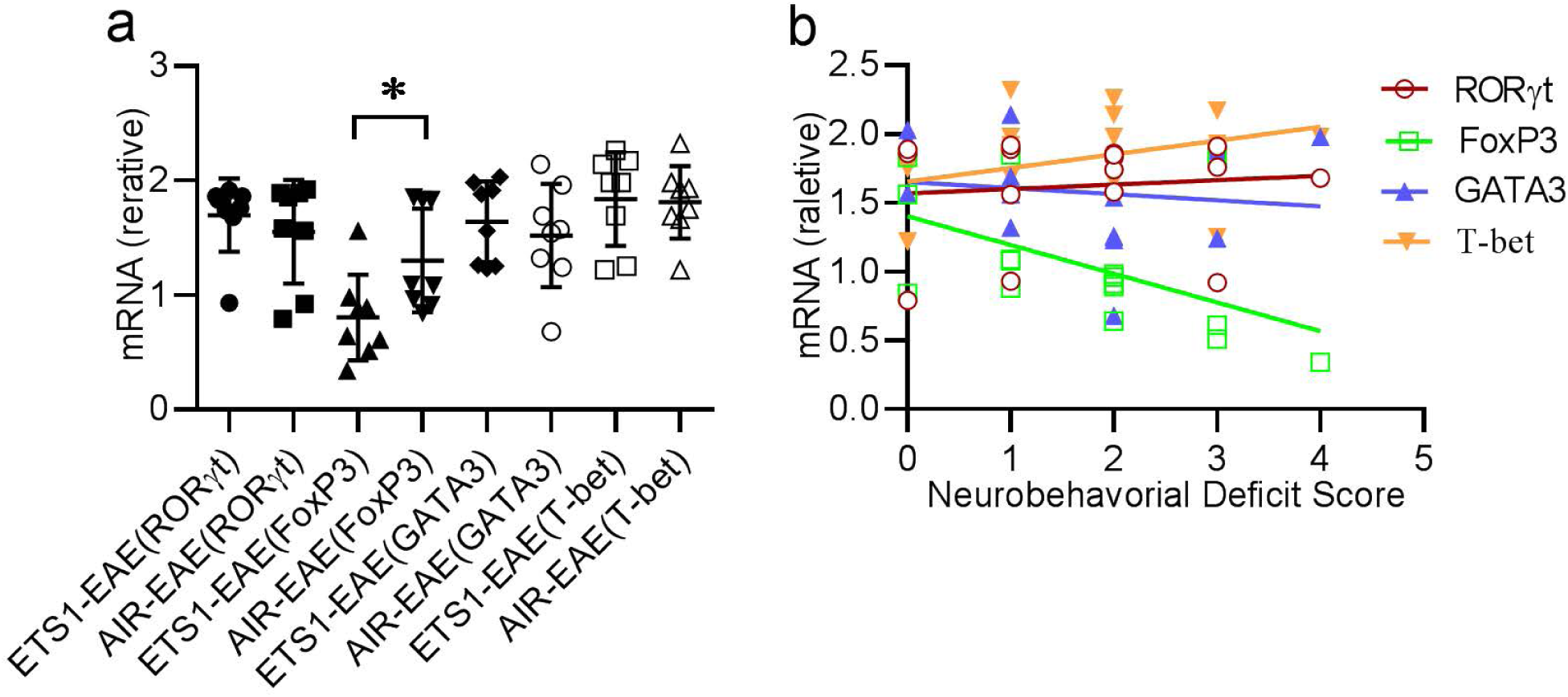
Effects of ETS exposure on the levels of transcription factors of CD4^+^cells and their correlation with neurobehavioral deficit scores. (a), On the 15 d after EAE induction, the brain and spinal cord were homogenized, and CD4^+^cells were purified by flow cytometry. Detection of RORγt, Foxp3, GATA3 and T-bet, mRNA expression in CD4^+^ cells was performed using real-time RT-PCR (n = 8). ETS1 exposure down-regulated Foxp3 expression in CD4^+^ cells. (b), Foxp3 expression level was negatively correlated with the neurobehavioral deficit score (R^2^ = 0.2747, r = −0.5241, n = 16). Data represent the mean ± SEM. \**P* < 0.05; Test used: Unpaired t-test analysis.

### Effects of early neonatal ETS exposure on inflammatory mediators in EAE rats

The imbalance between Treg cells and Teff cells affects the release of inflammatory mediators in the brain and spinal cord of EAE rats, and it may aggravate the pathogenesis of the disease. To this end, we further examined the cerebrospinal fluid (CSF) expression levels of IL-4, IL-10, IL-17, and INF-γ. In the CSF of ETS exposed rats, the level of IL-10, which was released by Treg cells, was down-regulated (Fig. 5b), while the level of IL-17, which promotes inflammation, was up-regulated (Fig. 5a). The levels of IL-4 and INF-γ were not significantly affected by ETS exposure (Fig. 5c, d, respectively).

**Fig 5.**
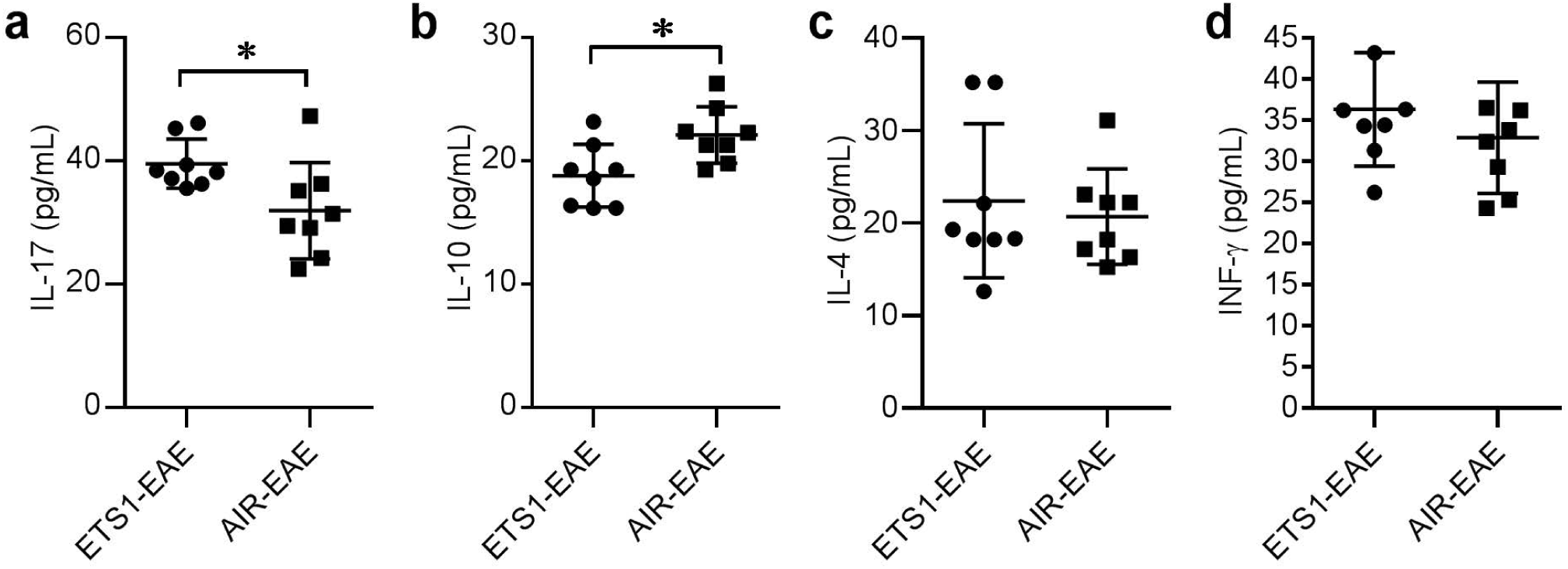
Early-life ETS exposure inhibits the expression of IL-10 in EAE rats. The concentrations of IL-17, IL-10, IL-4, and INF-γ were determined using ELISA 15 d after EAE induction. (a), Early-life ETS exposure up-regulated IL-17 concentration in the CSF of EAE rats (\**P* < 0.05, n = 8). (b), Early-life ETS exposure down-regulated IL-10 concentration in the CSF of EAE rats (\**P* < 0.05, n = 8). (c, d), Early-life ETS exposure did not affect the concentration of IL-4 (*P =* 0.6312) and INF-γ (*P* = 0.3317), respectively, in the CSF of EAE rats (n = 8). Data represent the mean ± SEM. Error bars indicate SEM. * *P* < 0.05; Test used: Unpaired t-test analysis

## Discussion

Early-life environmental factors exposure has a long-term impact on the development and function of the immune system, which ultimately affects the susceptibility and sensitivity to autoimmune diseases of the nervous system (Krementsov and Teuscher, 2013; Bar-Or et al., 2016; Pronovost and Hsiao, 2019). In this study, we found that neonatal ETS exposure increased the EAE severity but not the EAE incidence in adult rats. Furthermore, there is a critical time window period for neonatal ETS exposure to develop sensitivity to EAE. Evidence was provided to show that this sensitivity was related to the suppression of CD4 ^+^ CD25 ^+^ regulatory T cell differentiation. These data indicated that early-life exposure to ETS exacerbates MS severity in adulthood, but may not affect MS incidence.

The demyelination of the central nervous system is a characteristic pathological change of EAE / MS, which is related to the severity of the disease. We found that the demyelination area of rats exposed to ETS during P1 - P7 was more extensive by semi-quantitative analysis. In addition, our study showed that the axon density of the white matter area also decreased in the ETS1-EAE group rats, suggesting that the prognosis of the disease was worse. Axon damage is a pathological indicator for irreversible nervous system damage (Trapp et al., 1998; Preziosa et al., 2019).

The central infiltration level of CD4^+^ cells reflects the degree of immune activation during the onset of MS (Raphael et al., 2020). Our immunohistochemical staining showed that early-life ETS exposure promoted the infiltration of CD4^+^ cells in the spinal cord of EAE rats. Moreover, the results of the quantitative analysis of the absolute number of CD4+ cells isolated from brain and spinal cord tissues by flow cytometry also confirmed the results of CD4 immunohistochemical analysis. Further studies of lymphocytes from the brain and spinal cord showed that the number of CD4^+^ CD25^+^ Treg was smaller in the ETS1-EAE group, and the number of CD4^+^ CD25^−^ Teff cells was more abundant, but there was no difference found in peripheral spleen lymphocyte analysis.

Studies have shown that Treg cells account for more than 15% of CD4^+^ T cells in the rat cerebrum, and contribute to the immunosurveillance and immunomodulation in the cerebrum under steady-state (Xie et al., 2015). The individual gene expression levels of Treg cells in the brain are also significantly higher than that of peripheral Treg cells (Josefowicz et al., 2012). The difference in gene expression between central and peripheral Treg cells suggests that their function and origin pathway may be different. Indeed, early-life ETS exposure has a significant effect on the proliferation of central colonized Treg cells, but not the peripheral Treg cells, indicating a strong role of Treg in the development of the sensitivity to EAE. Foxp3 is the transcription factor that specifies the Treg cell lineage (Josefowicz et al., 2012). Given that the interaction between astrocytes and Treg cells contributes to the maintenance of Treg cell identity in the brain (Xie et al., 2015), the damage of central Treg cells by early-life ETS exposure may be indirectly affected by astrocytes dysfunction. Indeed, we found increased GFAP cells in all EAE groups. However, it is not clear whether astrocytes selectively contribute to any specific early time window exposure to ETS, which warrants further studies.

CD4 ^+^ CD25^-^ Teff cells are mainly composed of Th1, Th2, and Th17 cells (Axtell et al., 2010; Fletcher et al., 2010). The transcription factors related to the differentiation and activation of these cells are T-bet, GATA3, and RORγt, respectively (Stadhouders et al., 2018). Th1 and Th17 cells are considered to be the critical pathogenic CD4^+^T cell subsets leading to MS (Fletcher et al., 2010; Yadav et al., 2015). In the present study, Foxp3 expression level was significantly decreased in the ETS1-EAE group, which negatively correlated with the neurobehavioral deficit score. Transcription factors related to CD4 ^+^ CD25^-^ Teff cells were not dramatically changed and showed no significant correlation with the neurobehavioral deficit score. Therefore, early-life EST exposure decreases Foxp3 expression and suppresses Treg cells’ function.

To study the changes of the Treg and Teff cells associated cytokines levels, the levels of IL-10, IL-17, IL-4, and INF-γ in the CSF were measured at the EAE onset stage. We found that the level of IL-10 was down-regulated in ETS exposed rats, while IL-17 level was up-regulated, which was consistent with the down-regulation of Treg cells in central lymphocytes and the up-regulation of Teff cells. The results of the animal study are also similar to those found in the MS patients, where the level of IL-10 is down-regulated, and the levels of Th17 cells and IL-17 are up-regulated (Danikowski et al., 2017). It is believed that IL-10 secreted by Treg cells suppresses the Th1 and Th17 cells’ autoimmune activity. Therefore, it is plausible to reason that inhibition of Foxp3 expression in the brain and spinal cord by neonatal ETS exposure disrupts the Treg /Teff cells balance and leads to an aggravation of the pathogenesis of EAE.

In summary, although there is plenty of evidence that smoking is a risk factor for the recurrence and progression of MS in adulthood, the long-term effects of perinatal and postnatal ETS exposure on MS are not clear. Through animal experiments, we found for the first time that ETS exposure in the neonatal period did not alter the incidence of EAE, but increased the severity of the disease. Future research is warranted to examine the underlying mechanisms and their clinical applications.

## Acknowledgments

This work was supported by the Basic Public Welfare Research Plan of Zhejiang Province (LGF18H090018) to Z.W. Wang. Financial supports to Dr. S.T. Hou was from grants from the National Natural Science Foundation of China (81871026); Shenzhen-Hong Kong Institute of Brain Science-Shenzhen Fundamental Research Institutions (2019SHIBS0002); Shenzhen Science and Technology Innovation Committee Research Grants (JCYJ20180504165806229; KQJSCX20180322151111754). STH is also supported by the Guangdong Innovation Platform of Translational Research for Cerebrovascular Diseases and SUSTech-UQ Joint Center for Neuroscience and Neural Engineering (CNNE).

## Authors contributions

ZWW, LPW, FFZ, and CLW performed animal surgery, behavioral studies, data analysis. ZWW wrote the first draft of the manuscript. HST guided the manuscript writing and revised the final draft.

## Notes

### Competing Interest Statement

The authors have declared no competing interest.

